# Interactive Effects of Biological Maturation and Relative Age Effect on Talent Identification for U16 Elite Soccer Players

**DOI:** 10.64898/2026.04.02.716019

**Authors:** Yikang Gong, Weichao Jiang, Xiang Li, Yuan Li, Wei Zhang, Donghao Wang, Hua Wang, Chong Luo

**Affiliations:** Shanghai University of Sport, Shanghai, 200438, China; Beijing Sport University, Beijing, 100084, China

**Keywords:** Biological Maturation, Relative Age Effect, Talent Selection

## Abstract

This retrospective study aims to explore the interactive effects of biological maturation and relative age effect (RAE) on talent identification. 56 male elite soccer players matched for chronological age (15.08±0.41 years) were studied. Test items included anthropometry (height, body mass, sitting height, leg length, BMI and Quetelet index), physiology (power, speed, agility, speed endurance and aerobic performance), soccer-specific skills (passing, shooting and dribbling), psychology (achievement motivation, orientation and resilience) and biological maturation (APHV) tests. The test results were analyzed independent sample t-test, Pearson correlation analysis, and stratified regression. Conclusion: Biological maturation significantly influences anthropometry (height, weight and Quetelet index), lower limb explosive, and speed (single-leg jump, standing triple jump, and 30-m sprint) in U16 male elite soccer players in Shanghai. The relative age effect shows no significant impact on talent selection indicators, which is attributed to the accumulated training load effect. The mechanisms of biological maturation and RAE in youth soccer talent selection are distinct and operate independently.

## 1 Introduction

Interindividual variations play a critical role in the development of youth soccer players, particularly concerning biological maturation and the relative age effect. Biological maturation refers to the progressive development of individual organs, systems, and tissues, and is commonly described in terms of three dimensions: maturity status, maturity timing and maturity tempo ^[1],[2]^. Maturity status denotes the individual’s current stage of development (i.e. pre-puberty, puberty, and post-puberty), while maturity timing indicates the age at which specific maturity events occur, such as the growth spurt or menarche ^[1]^. Maturity tempo, on the other hand, reflects the rate at which this developmental progression, and is marked by considerable interindividual variability in pubertal timing ^[2]^. Current study suggests that pubertal development is regulated by the interaction of genetic predispositions and environmental influences through the hypothalamic-pituitary-gonadal axis ^[3]^. These complex interactions drive variations in pubertal timing, resulting in early, normal, or late maturation ^[3]^. Variations in maturity status confer distinct physiological and morphological advantages to early maturing players, such as advanced skeletal age, greater muscle hypertrophy, and superior performance in endurance, power and speed capacities ^[4]^. These early physical advantages often lead coaches to preferentially select early maturing players, granting them increased playing time and a greater likelihood of selection for elite teams ^[5]^.

Relative Age (RA) denotes the age differences between individuals within the same chronological age group ^[6]^. The Relative Age Effect (RAE), also known as the birthdate effect, describes the phenomenon whereby an athlete’s performance and likelihood of selection are influenced by their birthdate relative to established age-group cutoff dates ^[7]^. In youth soccer, players are typically grouped by chronological age cohorts. This grouping results in age differences of up to nearly 12 months between players born early (e.g., January) and late (e.g., December) within the same selection year. Due to this disparity, players born earlier in the year often exhibit superior physical fitness compared to their younger peers and are more likely to be perceived as more skilled or talented. Consequently, talent identification and selection processes tend to disproportionately favor early born players, potentially overlooking late-born players who may possess comparable or even superior long-term potential ^[8]^.

Previous study has frequentl<u>y</u> considered biological maturation as a key factor contributing to RAE ^[9]^. However, recent studies have shown that biological maturation and RAE should not be viewed merely as subordinate concepts, but rather as distinct and independent constructs ^[5],[10]^. Mechanistically, individual differences in biological maturation are primarily influenced by the interaction of genetic, environmental, and behavioral factors, such as chronic malnutrition, illness, and climate conditions ^[11]^. In contrast, the RAE is primarily generated by institutional practices, particularly age-based grouping systems, which create performance differences between players born earlier and later within the same selection year ^[12]^. In terms of scope, biological maturation can have a maximum effect of 5-6 years in a single age group ^[13], [14], [15]^, while the RAE has a maximum impact of 12 months in a single age group.

Teaching youth soccer talent identification is the process of identifying or selecting existing participants with the potential to become elite players ^[1]^. Talent selection generally involves evaluating anthropometry (height, body mass, sitting height, leg length, skinfolds, BMI and Quetelet index), physiology (power, speed, agility, speed endurance and aerobic performance), soccer-specific skills (passing, shooting and dribbling), and psychology (achievement motivation, orientation and resilience). Multiple studies have demonstrated that biological maturation and RA significantly affect anthropometry, physiology, soccer-specific skills, and psychology in youth soccer talents ^[1],[2],[5],[6]^. The study treats biological maturation and RA as two independent variables to investigate the interactive effects of biological maturation and the RAE on the physiology, soccer-specific skills, and psychology of U16 elite soccer players. Thereby, it aims to refine the theoretical framework of soccer talent development and enhance the quality and efficiency of talent selection.

## 2 Methods

This study calculated the sample size using G*Power, selecting 56 U16 youth football players from the youth academies of Shanghai Port Football Club and Shanghai Shenhua F.C. Participants were recruited for this study between 01/03/2026 and 08/03/2026. All research participants provided written informed consent. Before all tests, each participant administered a psychological questionnaire under the supervision of the test personnel, who ensured the completion of the entire questionnaire. Soccer-specific skills tests (passing, shooting and dribbling) were conducted in the morning of the test day, while physiological tests (power, speed, agility, speed endurance and aerobic performance) were scheduled in the afternoon. Before the commencement of all tests, all participants were led by the test personnel through a 20-minute warm-up session on the soccer field. Participants were then randomly divided into five groups. Five test personnel were deployed to each test location by the requirements to conduct the testing and collect the data. Two test personnel recorded the results for each test, and a demonstration was conducted before each test. Participants were allowed to perform a trial run before the formal test. Except for height, sitting height, weight, and countermovement jump (CMJ), all other tests were conducted on the soccer field.

### 2.1 Anthropometry, Biological Maturation, and Relative Age Effect

*Height* was measured using an anthropometer, with participants barefoot and standing upright on the base plate of the instrument. The heels, sacral region, and scapular areas were tightly aligned with the upright column of the anthropometer. The measurer stood on the side, adjusted the participant’s posture to ensure the eyes were directly forward, and raised the horizontal headpiece to the top of the head to record the height in centimeters (cm), accurate to one decimal place.

*Sitting height* was measured using the same anthropometer, with participants seated on the sitting board and maintaining an upright torso. The spinal column and sacral region were aligned vertically between the shoulders and sacrum, without leaning against the wall. The head was positioned at the eye-ear plane. The sitting height was recorded in centimeters (cm), accurate to one decimal place.

*Leg length* was calculated by subtracting sitting height from stature.

*Body weight* was measured using the KISTLER multi-component force plate, with all participants wearing competition attire and maintaining an upright posture during testing. The weight was recorded in kilograms (kg), accurate to two decimal places.

*BMI and Quetelet index* were expressed as ratios to meet the requirements of the maturity offset formula. BMI = weight (kg) / height (m) ^ 2; Quetelet index = weight (kg) / height (cm) * 1000.

#### Biological Maturation Assessment

This study employed a non-invasive method to evaluate biological maturation, using a prediction equation of maturity offset constructed based on the Shanghai Longitudinal Growth and Development Study (SLGDS) in 2024 ^[18]^. The maturity offset (MO) for male players was calculated using the formula: MO = -15.553 + 0.705 × age + 0.067 × sitting height + 0.063 × BMI. CA was determined by subtracting the player’s birth date from the testing date, with the difference calculated using the DAYS360 function in Excel. Age at peak height velocity (APHV) was estimated as the difference between CA and MO. Maturity was estimated using Z-scores, and the mean and standard deviation of APHV were obtained from the literature review ^[19]^. Z-scores were used to categorize players into maturational status (early, normal, or late maturity) ^[11], [20], [21]^, with a Z-score of -0.5 to +0.5 defining normal maturity, a Z-score > +0.5 classified as late maturity, and a Z-score < -0.5 classified as early maturity ^[22]^.

#### Relative Age Assessment

The players’ birth dates were categorized into four quarters. Referring to the study by Parr et al. ^[13]^, the relative age was converted into a continuous variable by dividing the number of days between the player’s birth date and the selection cutoff date by 365 ^[23]^.

### 2.2 Physical performance evaluation

Physical performance evaluation primarily focuses on three dimensions: power, agility and speed. Each test was performed twice, and the best result was selected for analysis. Power was evaluated using three tests: CMJ, single-leg jump (SJ), and standing triple jump (STJ). Agility was measured with the Illinois Agility Test (IAT) and the 505 Agility Test, both of which utilize electronic timing systems. Speed was assessed through three sprint tests: 15-m, 30-m, and 5×25-m sprints (see Table 1).

**Table 1.** Physical performance tests.

| Test Items |  | Test Tools | Test Method | Measurement Unit | Test Times | Interval Time |
| --- | --- | --- | --- | --- | --- | --- |
| Power Tests | CMJ | Kistler 3D force platform | CMJ is a vertical jump that involves a downward movement (countermovement) before an upward leap. | Height (m) | 2 | 1-2 minutes |
|  | SJ | Tape measure | This exercise involves jumping off one leg as high as possible while maintaining balance and control. Distance measurement is the shortest distance from the heel to the jump line. | Distance (cm) | Twice on each foot | 1-2 minutes |
|  | STJ | Tape measure | The player must make sure they land their hop with the same foot that they took off with, then they must alternate to the other foot to land from the step. They then must make the jump from this foot and can land on both feet. The distance measurement is the shortest distance from the heel to the jump line. | Distance (cm) | 2 | 1 minute |
| Agility Tests | IAT | Smart Speed timing system | Players perform a standing start, following a designated course, with the total time recorded from the start to the finish line. | Time (s) | 2 | 1-2 minutes |
|  | 505 Agility Test | Smart Speed timing system | Players perform a standing start, sprinting 5 m to the turn line and then turning 180°, with the total time recorded from the start to the finish line. | Time (s) | 2 | 1 minute |
| Speed Tests | 15-m/30-m sprints | Smart Speed timing system | Players perform a standing start, initiating their sprint 30 cm behind the timing gate, and accelerating to the finish line cone. | Time (s) | 2 | 2 minutes |
|  | 5×25-m sprints | Smart Speed timing system | Players perform a standing start, sequentially turning back and knocking down markers at 5 m, 10 m, 15 m, 20 m, and 25 m intervals, with the total time recorded from the start to the finish line." | Time (s) | 1 |  |

CMJ=Countermovement Jump, SJ=Single-leg Jump, STJ=Standing Triple Jump, IAT=Illinois Agility Test.

**Table 2.** Differences in testing indicators between early maturing and normal maturing players.

| Indexes | Grouping (Mean ± Standard Deviation) | | t | p | Difference and 95%CI |
| --- | --- | --- | --- | --- | --- |
|  | Early Maturity (n=34) | Normal Maturity (n=22) |  |  |  |
| Height (cm) | 176.87±5.34 | 169.86±4.49 | 4.190 | **0.000\*\*** | 7.01 (3.62-10.40) |
| Weight (kg) | 68.15±6.42 | 56.02±6.49 | 5.832 | **0.000\*\*** | 12.13 (7.92-16.35) |
| Sitting Height (cm) | 94.98±2.42 | 90.09±3.66 | 5.110 | **0.000\*\*** | 4.90 (2.95-6.84) |
| BMI | 21.80±2.03 | 19.38±1.84 | 3.841 | **0.000\*\*** | 2.42 (1.14-3.70) |
| Quetelet Index | 385.30±34.35 | 329.39±33.73 | 5.071 | **0.000\*\*** | 55.9 (33.62-78.19) |
| Leg Length (cm) | 81.89±4.18 | 79.77±3.13 | 1.732 | 0.090 | 1.14 (-0.36-4.60) |
| CMJ (Newton) | 43.58±4.49 | 41.18±3.66 | 1.783 | 0.083 | 2.40(-0.33 - 5.13) |
| SJL (cm) | 211.21±16.94 | 200.39±7.33 | 2.401 | **0.021\*** | 10.82(1.70 - 19.95) |
| SJR (cm) | 211.29±14.96 | 202.73±12.69 | 1.882 | 0.068 | 8.57(-0.65 - 17.78) |
| STJ (m) | 6.96±0.49 | 6.63±0.47 | 2.101 | **0.043\*** | 0.33(0.01 - 0.64) |
| IAT (sec) | 15.98±0.71 | 16.05±0.51 | -0.342 | 0.737 | -0.07(-0.49 - 0.35) |
| 505 Agility Test (sec) | 2.40±0.12 | 2.40±0.11 | -0.131 | 0.897 | -0.00(-0.08 - 0.07) |
| 15-m Sprint (sec) | 2.23±0.19 | 2.37±0.15 | -2.431 | **0.020\*** | -0.14(-0.25 - -0.02) |
| 30-m Sprint (sec) | 4.12±0.26 | 4.40±0.23 | -3.382 | **0.002\*** | -0.27(-0.44 - -0.11) |
| 5×25 Sprints (sec) | 33.97±2.20 | 34.47±1.46 | -0.801 | 0.429 | -0.50(-1.77 - 0.77) |
| Spot Passing (score) | 3.75±2.21 | 3.13±2.09 | 0.894 | 0.377 | 0.625(-0.790-2.040) |
| Shooting (score) | 4.46±2.17 | 4.31±1.99 | 0.215 | 0.831 | 0.146(-1.225-1.517) |
| Dribbling (sec) | 10.43±0.69 | 10.39±0.42 | 0.248 | 0.805 | 0.048(-0.341-0.437) |
| AM-S | 4.25±0.54 | 4.30±0.60 | -0.252 | 0.802 | -0.046(-0.414-0.322) |
| AM-F | 2.23±0.57 | 2.47±1.00 | -0.964 | 0.341 | -0.240(-0.743-0.263) |
| AM | 2.02±0.86 | 1.83±1.37 | 0.505 | 0.586 | 0.194(-0.520-0.908) |
| T | 30.29±3.65 | 30.38±4.18 | -0.067 | 0.947 | -0.083(-2.611-2.444) |
| E | 24.04±3.84 | 23.94±3.87 | 0.084 | 0.934 | 0.104(-2.413-2.621) |
| TEOSQ | 54.33±6.97 | 54.31±7.26 | 0.009 | 0.993 | 0.021(-4.610-4.652) |
| F1 | 20.25±2.85 | 21.50±3.23 | -1.290 | 0.205 | -1.250(-3.212-0.712) |
| F2 | -3.50±3.02 | -4.06±4.06 | 0.503 | 0.618 | 0.563(-1.703-2.828) |
| F3 | 16.92±2.96 | 16.88±2.53 | 0.046 | 0.963 | 0.042(-1.787-1.870) |
| F4 | 6.00±3.34 | 5.88±4.81 | 0.097 | 0.923 | 0.125(-2.479-2.729) |
| F5 | -1.25±5.24 | -2.37±4.33 | 0.711 | 0.481 | 1.125(-2.076-4.326) |
| F | 38.42±11.77 | 37.81±14.42 | 0.145 | 0.885 | 0.604(-7.814-9.022) |

\*, p<0.05; \*\*, p<0.01; \*\*\*, p<0.001.
CMJ=Countermovement Jump, SJL=Single-leg Jump (Left), SJR=Single-leg Jump (Right), STJ=Standing Triple Jump, IAT=Illinois Agility Test, AM-S=Achievement Motivation (Approach Success), AM-F=Achievement Motivation (Avoidance Failure), AM=Achievement Motivation, T=Task Orientation, E=Ego Orientation, TEOSQ=Task and Ego Orientation in Sports Questionnaire, F1=Goal-Oriented, F2=Emotional Control, F3=Positive Cognition, F4=Family Support, F5=Interpersonal Assistants, F=Youth Psychological Resilience Scale.

**Table 3.** Performance metrics differences across different relative age groups.

| Indexes | Grouping (Mean ± Standard Deviation) | | | | f | p |
| --- | --- | --- | --- | --- | --- | --- |
|  | Q1 | Q2 | Q3 | Q4 |  |  |
| Height | 174.01±4.75 | 174.65±6.04 | 174.97±8.07 | 171.87±6.6 | 0.379 | >0.05 |
| Weight | 62.68±7.89 | 62.87±10.31 | 62.48±8.28 | 62.37±9.09 | 0.198 | >0.05 |
| Sitting Height | 92.98±7.89 | 93.68±5.21 | 92.66±2.93 | 92.34±3.16 | 0.22 | >0.05 |
| BMI | 20.66±1.97 | 21.18±2.71 | 20.37±2.01 | 21.07±2.49 | 0.254 | >0.05 |
| Quetelet Index | 359.69±38.93 | 370.49±52.57 | 356.48±38.38 | 362.31±40.32 | 0.203 | >0.05 |
| Leg Length | 81.03±2.87 | 80.97±2.78 | 82.32±5.94 | 79.53±4.15 | 0.66 | >0.05 |
| CMJ | 42.30±4.15 | 41.89±3.79 | 44.44±5.31 | 42.12±4.28 | 0.379 | >0.05 |
| SJL | 209.27±10.72 | 197.47±11.44 | 216.89±19.62 | 207.71±10.13 | 0.379 | >0.05 |
| SJR | 209.36±9.55 | 202.51±14.74 | 214.56±20.22 | 206.86±10.56 | 0.379 | >0.05 |
| STJ | 6.89±0.43 | 6.70±0.53 | 6.96±0.58 | 6.79±0.50 | 0.379 | >0.05 |
| IAT | 15.91±0.38 | 15.90±0.56 | 16.14±0.97 | 16.22±0.58 | 0.379 | >0.05 |
| 505 Agility Test | 2.44±0.10 | 2.38±0.09 | 2.39±0.16 | 2.36±0.11 | 0.379 | >0.05 |
| 15-m Sprint | 2.32±0.22 | 2.28±0.15 | 2.28±0.24 | 2.26±0.14 | 0.379 | >0.05 |
| 30-m Sprint | 4.27±0.31 | 4.21±0.24 | 4.23±0.37 | 4.22±0.22 | 0.379 | >0.05 |
| 5×25 Sprints | 33.91±1.60 | 33.84±1.68 | 33.97±2.22 | 35.48±2.32 | 0.379 | >0.05 |
| Spot Passing | 3.45±2.38 | 3.85±2.67 | 2.78±1.56 | 3.86±1.46 | 0.492 | >0.05 |
| Shooting | 4.36±1.63 | 4.23±2.01 | 5.11±2.67 | 3.86±2.19 | 0.523 | >0.05 |
| Dribbling | 10.34±0.48 | 10.33±0.43 | 10.65±0.95 | 10.40±0.42 | 0.618 | >0.05 |
| AM-S | 4.31±0.50 | 4.22±0.70 | 4.24±0.56 | 4.33±0.44 | 0.087 | >0.05 |
| AM-F | 2.13±0.86 | 2.35±0.86 | 2.53±0.70 | 2.31±0.59 | 0.449 | >0.05 |
| AM | 2.19±1.25 | 1.87±1.22 | 1.71±0.93 | 2.02±0.83 | 0.342 | >0.05 |
| T | 30.09±2.91 | 29.31±4.94 | 30.78±3.23 | 32.00±3.46 | 0.800 | >0.05 |
| E | 23.64±4.13 | 23.23±3.79 | 24.56±4.45 | 25.29±2.56 | 0.522 | >0.05 |
| TEOSQ | 53.75±6.56 | 52.54±7.99 | 55.33±7.07 | 57.29±5.7 | 0.776 | >0.05 |
| F1 | 21.45±2.25 | 20.31±3.40 | 21.00±3.08 | 20.14±3.67 | 0.386 | >0.05 |
| F2 | -3.27±2.87 | -3.54±4.48 | -5.11±3.02 | -3.00±2.58 | 0.651 | >0.05 |
| F3 | 17.91±1.64 | 16.54±2.96 | 16.89±2.03 | 16.00±4.36 | 0.799 | >0.05 |
| F4 | 6.09±2.98 | 5.38±4.70 | 7.33±1.87 | 5.00±5.69 | 0.582 | >0.05 |
| F5 | -0.64±3.38 | -3.54±5.52 | -1.11±4.78 | -0.71±5.71 | 0.927 | >0.05 |
| F | 41.55±10.18 | 35.15±15.70 | 39.00±8.35 | 37.43±15.84 | 0.502 | >0.05 |

\*, $p < 0.05$ ; \*\*, $p < 0.01$ ; \*\*\*, $p < 0.001$ .
CMJ=Countermovement Jump, SJL=Single-leg Jump (Left), SJR=Single-leg Jump (Right), STJ=Standing Triple Jump, IAT=Illinois Agility Test, AM-S=Achievement Motivation (Approach Success), AM-F=Achievement Motivation (Avoidance Failure), AM=Achievement Motivation, T=Task Orientation, E=Ego Orientation, TEOSQ=Task and Ego Orientation in Sports Questionnaire, F1=Goal-Oriented, F2=Emotional Control, F3=Positive Cognition, F4=Family Support, F5=Interpersonal Assistants, F=Youth Psychological Resilience Scale.

### 2.3 Soccer-specific Skills

#### Spot passing test

Players start by dribbling the ball from behind the marker line into the passing zone. They then alternate using their left and right feet to sequentially pass the ball to points A, B, and C. After each pass, the player returns to the starting line. Scoring is determined based on the ball’s landing position: 1 point for landing within the outer circle and 2 points for landing within the inner circle (see Figure 1-A).

**Figure 1.**
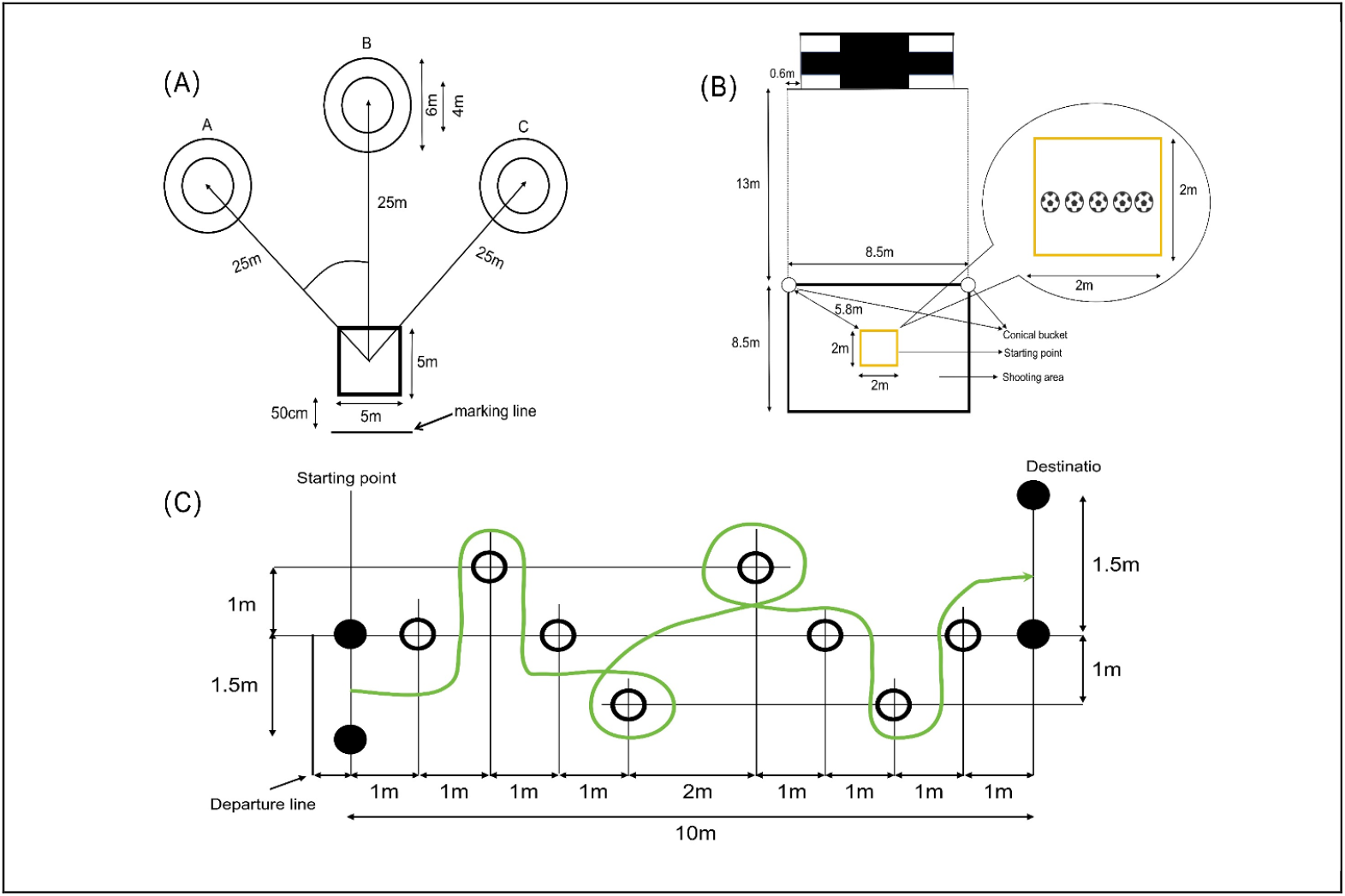
Soccer-specific skills test (A) spot passing test (B) shooting test (C) dribbling test

**Figure 2.**
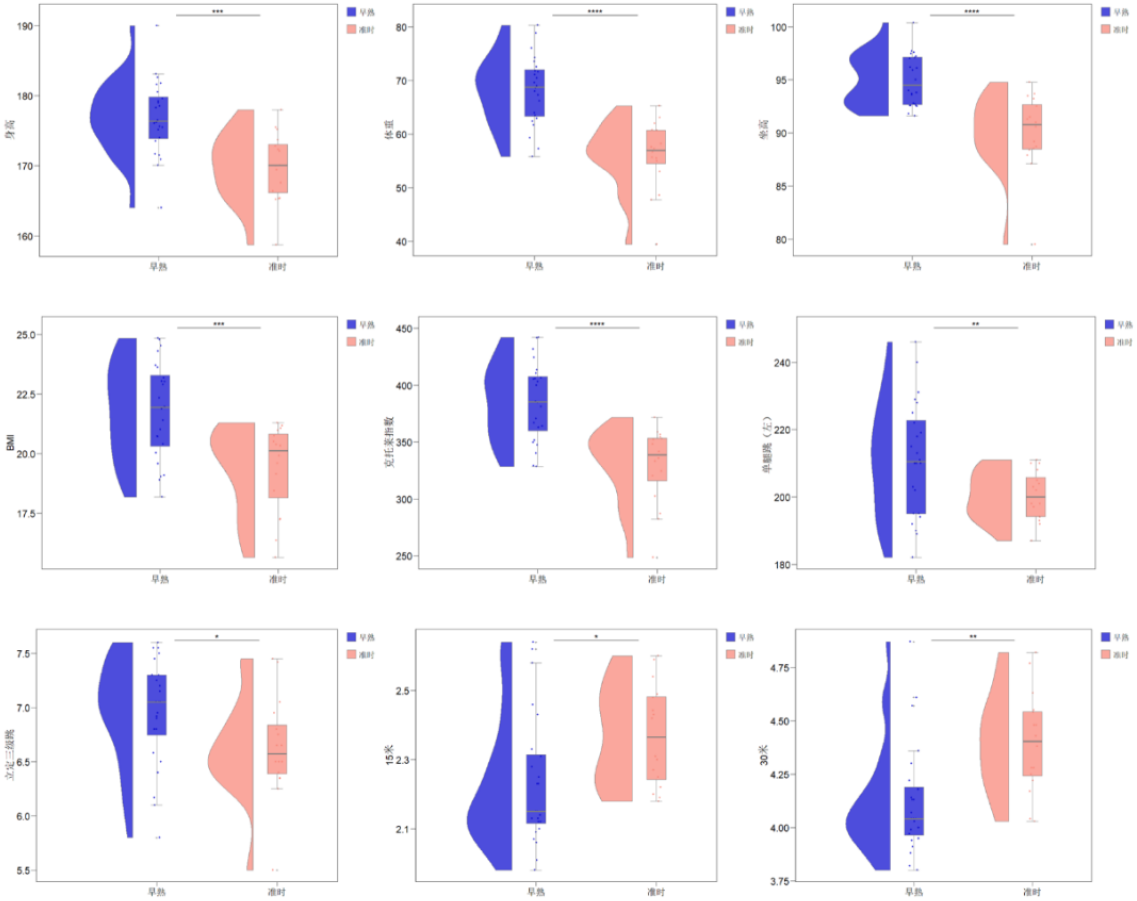
Significance differences in performance metrics across maturational groups

**Figure 3.**
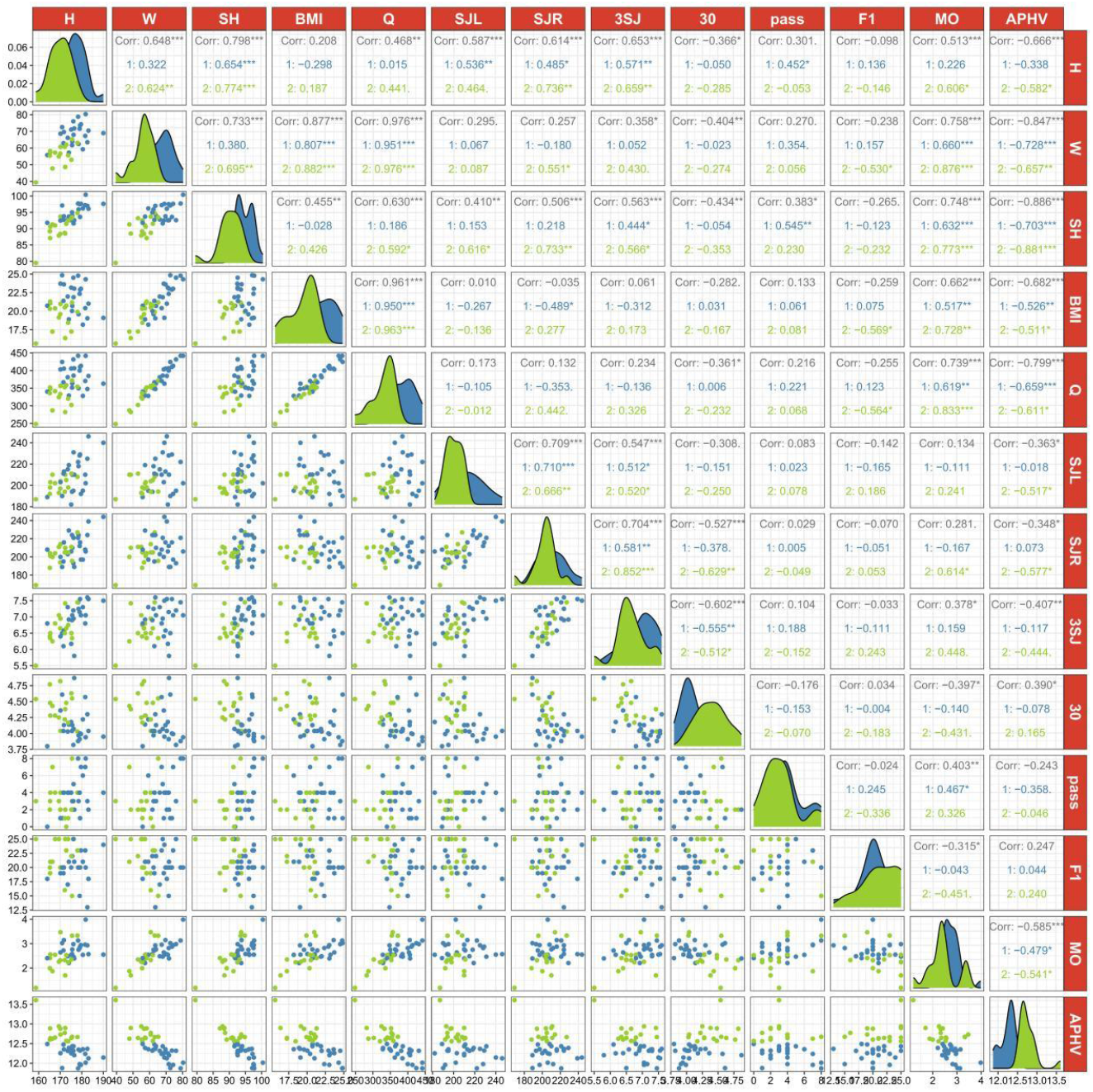
Scatter plot matrix for significant correlation testing of the variables. H=Height, W= Weight, SH=Sitting Height, Q=Quetelet Index, SJL=Single-leg Jump (Left), SJR=Single-leg Jump (Right), 3SJ =Standing Triple Jump, 30=30-m Sprint, Pass=Spot Passing, F1=Goal-Oriented, MO=Maturity Offset, APHV=Age at Peak Height Velocity.

#### Shooting test

Players start from the starting line facing away from the goal, turn around, and then shoot 5 times with each foot in the goal area. After each shot, the player must knock down a cone and wait until the ball crosses the line before proceeding. Scoring criteria: 3 points for scoring in the top corner, 1 point for the bottom corner, and 0 points for missed shots. The total score is the sum of shot scores (see Figure 1-B).

#### Dribbling test

Upon hearing the signal, the athletes followed the marked route, navigating around the cones to reach the finish line with the timer stopping when the athlete’s body crosses the line. Two trials are conducted with a 1 – 2-minute rest period between them, and the best performance is recorded as the result (see Figure 1-C).

### 2.4 Psychology

The Sport Achievement Motivation Scale (SAMS) was used to assess participants’ motivation. The scale consists of two subscales: approach success and avoidance failure, with total scores calculated as the difference between the two subscales (higher scores indicate stronger motivation). The reliability coefficients of the scale ranged from 0.894 to 0.923 ^[28]^. The Task and Ego Orientation in Sports Questionnaire (TEOSQ) was designed to measure goal orientation in sports, comprising two subscales: task orientation and ego orientation. The total score is calculated by summing the scores of the two subscales, with higher scores indicating a stronger tendency toward the respective orientation. The reliability coefficients of the scale ranged from 0.781 to 0.876 ^[28]^.

Psychological resilience was measured using the Youth Psychological Resilience Scale (YPRS) ^[17]^. The 27-item scale employs the Likert 5-Point Scale for responses ^[16]^. The test-retest reliability coefficient was 0.83, and the internal consistency coefficient was 0.92 ^[17]^.

### 2.5 Statistics analysis

The study utilized SPSS 26.0 for data analysis on anthropometry, physiology, soccer-specific skills and psychology, biological maturation and RA, with a significance level set at *P* < 0.04. Normality of the data was assessed using the Shapiro-Wilk test. Comparative analyses were conducted using t-tests and one-way ANOVA. Correlation analysis was employed to examine the relationships between talent selection indicators and biological maturation and RA. The hierarchical regression model was implemented to investigate the main effects and the interaction effects of biological maturation and RA. The regression analysis was conducted in three hierarchical models: Model 1 included chronological age as a covariate; Model 2 added the main effects of biological maturation and RA; and Model 3 introduced the interaction effects between biological maturity and RA.

## 3 Results

Among the 56 U16 elite male soccer players, 34 (60%) were classified as early maturity, 22 (40%) as normal maturity, and no late maturity. A higher proportion of players born in the first two quarters of the year (Q1 = 16 and Q2 = 18, totaling 34 players, 60%) was observed compared to those born in the last two quarters (Q3 = 12 and Q4 = 10, totaling 22 players, 40%).

### 3.1 Variance Analysis

To assess the differences in performance metrics between early maturity and normal maturity, an independent sample t-test was conducted. The results showed that early maturing players exhibited significantly higher mean values in anthropometry compared to normal maturing players. Specifically, the average height of early maturing players was 176.87 ± 5.34 cm, compared to 169.86 ± 4.49 cm for average maturing players, with a difference of 7.01 cm (95% CI: 3.62 – 10.40). The weight difference was 12.13 kg (95% CI: 7.92 – 16.35), and the sitting height difference was 4.9 cm (95% CI: 2.95 – 6.84). The BMI difference was 2.42 (95% CI: 1.14 – 3.70), and the Quetelet index difference was 55.9 (95% CI: 33.62– 78.19).

In terms of physiology, the SJL, STJ, 15-m and 30-m sprints showed significant differences between the two groups (*p* < 0.05). The SJL for early maturing players was 211.21 ± 16.94 cm, compared to 200.39 ± 7.33 cm for normal maturing players, with a difference of 10.82 cm (95% CI: 1.70 – 19.95). The STJ for early maturing players was 6.96 ± 0.49 m, compared to 6.63 ± 0.47 m for normal maturing players, with a difference of 0.33 m (95% CI: 0.01 – 0.64). The 15-m sprint time for early maturing players was 2.23 ± 0.19 s, compared to 2.37 ± 0.15 s for normal maturing players, with a difference of -0.14 (95% CI: -0.25 – 0.02). The 30-m sprint time for early maturing players was 4.12 ± 0.26 s, compared to 4.40 ± 0.23 s for normal maturing players, with a difference of -0.27 (95% CI: -0.44 – 0.11). However, the CMJ, the IAT and the 505 Agility Test and 5×25 m sprints did not show significant differences between the two groups (*p* > 0.05), indicating that biological maturation did not significantly affect the agility and anaerobic endurance of U16 elite male soccer players.

Additionally, no significant differences were observed in soccer-specific skill performance (spot passing, shooting, and dribbling) and psychological indicators between the early and normal maturing groups (*p* > 0.05). Although early maturing players showed slightly higher mean scores in passing and shooting, the 95% confidence intervals for both differences included 0, indicating that the differences were not statistically significant.

A one-way ANOVA was conducted with birth quarter as the grouping variable to analyze the differences in performance metrics. Results revealed that the birth quarter did not significantly affect the talent identification metrics among U16 elite male soccer players in Shanghai (*p* > 0.05).

### 3.2 Correlation Analysis

Pearson correlation analysis was conducted to examine the relationships between various variables and biological maturation and RA, and the correlation coefficient matrix was generated using the R Programming Language. The maturity index used MO and APHV for analysis, and relative age was converted into a continuous variable ^[24]^. It can be observed that for male soccer players, MO and APHV were significantly correlated with height (r = 0.513, -0.666, *p* < 0.05), weight (r = 0.758, -0.847, *p* < 0.05), sitting height (r = 0.748, -0.886, *p* < 0.05), BMI (r = 0.662, -0.682, *p* < 0.05) and Quetelet index (r = 0.739-0.799, *p* < 0.05). However, leg length (r = 0.083, -0.192, *p* > 0.05) did not show significant correlation. The MO was significantly correlated with STJ (r = 0.378, *p* < 0.05) and 30-m sprint (r = -0.397, *p* < 0.05). APHV was significantly correlated with SJR (r = -0.363, *p* < 0.05), SJL (r = -0.348, *p* < 0.05), STJ (r = -0.407, *p* < 0.05), and 30-m sprint (r = 0.39, *p* < 0.05) (see Table 4). In terms of soccer-specific skills, the maturity index was positively correlated with passing (r = 0.403). In terms of psychology, maturity index was significantly correlated with target-oriented focus (r = -0.315). RA showed no significant correlation with any of the talent selection indicators.

**Table 4.** Correlation analysis between biological maturation, RAE and performance metrics.

| Indexes | MO | APHV | RA |
| --- | --- | --- | --- |
| Height | **0.513<sup>\*\*\*</sup>** | **-0.666<sup>\*\*\*</sup>** | 0.028 |
| Weight | **0.758<sup>\*\*\*</sup>** | **-0.847<sup>\*\*\*</sup>** | -0.081 |
| Sitting Height | **0.748<sup>\*\*\*</sup>** | **-0.886<sup>\*\*\*</sup>** | -0.033 |
| BMI | **0.662<sup>\*\*\*</sup>** | **-0.682<sup>\*\*\*</sup>** | -0.125 |
| Quetelet Index | **0.739<sup>\*\*\*</sup>** | **-0.799<sup>\*\*\*</sup>** | -0.102 |
| Leg Length | 0.083 | -0.192 | 0.078 |
| CMJ | 0.241 | -0.283 | -0.081 |
| SJL | 0.134 | **-0.363<sup>\*</sup>** | 0.117 |
| SJR | 0.281 | **-0.348<sup>\*</sup>** | -0.024 |
| STJ | **0.378<sup>\*</sup>** | **-0.407<sup>\*\*</sup>** | -0.016 |
| IAT | -0.068 | 0.066 | -0.011 |
| 505 Agility Test | -0.069 | 0.008 | 0.155 |
| 15-m Sprint | **-0.305<sup>\*</sup>** | 0.216 | 0.073 |
| 30-m Sprint | **-0.397<sup>\*</sup>** | **0.390<sup>\*</sup>** | 0.055 |
| 5×25 Sprints | -0.019 | 0.079 | -0.059 |
| Spot Passing | **0.403<sup>\*</sup>** | -0.329 | 0.001 |
| Shooting | 0.031 | -0.011 | 0.007 |
| Dribbling | 0.006 | 0.013 | 0 |
| AM-S | -0.262 | 0.091 | -0.069 |
| AM-F | 0.07 | 0.091 | -0.111 |
| AM | -0.184 | -0.018 | 0.043 |
| T | -0.282 | 0.112 | 0.196 |
| E | -0.03 | -0.072 | 0.102 |
| TEOSQ | -0.17 | 0.022 | 0.163 |
| F1 | **-0.315<sup>\*</sup>** | 0.247 | 0.141 |
| F2 | 0.04 | -0.013 | -0.16 |
| F3 | -0.095 | 0.08 | 0.179 |
| F4 | -0.241 | 0.107 | 0.075 |
| F5 | -0.032 | -0.097 | 0.208 |
| F | -0.172 | 0.068 | 0.132 |

\*, p<0.05; \*\*, p<0.01; \*\*\*, p<0.001;
MO=Maturity Offset, APHV=Age at Peak Height Velocity, RA=Relative Age, CMJ=Countermovement Jump, SJL=Single-leg Jump (Left), SJR=Single-leg Jump (Right), STJ=Standing Triple Jump, IAT=Illinois Agility Test, AM-S=Achievement Motivation (Approach Success), AM-F=Achievement Motivation (Avoidance Failure), AM=Achievement Motivation, T=Task Orientation, E=Ego Orientation, TEOSQ=Task and Ego Orientation in Sports Questionnaire, F1=Goal-Oriented, F2=Emotional Control, F3=Positive Cognition, F4=Family Support, F5=Interpersonal Assistants, F=Youth Psychological Resilience Scale.

**Table 5.** Hierarchical Regression Model of Biological Maturation and RAE on Performance Metrics.

| Indexes | Model 1 |  | Model 2 |  |  |  | Model 3 |  |  |  |  |
| --- | --- | --- | --- | --- | --- | --- | --- | --- | --- | --- | --- |
| | $\beta_1$ | $R^2$ | $\beta_1$ | $\beta_2$ | $\beta_3$ | $\Delta R^2$ | $\beta_1$ | $\beta_2$ | $\beta_3$ | $\beta_4$ | $\Delta R^2$ |
| Height | 0.074 | 0.006 | **-0.671\*\*\*** | 0.078 | **1.014\*\*\*** | 0.466 | **-0.676\*\*\*** | 0.375 | **1.179\*** | -0.328 | 0.002 |
| Weight | 0.224 | 0.05 | **-0.738\*\*\*** | -0.012 | **1.302\*\*\*** | 0.773 | **-0.724\*\*\*** | -0.785 | **0.874\*** | 0.853 | 0.009 |
| Sitting Height | 0.18 | 0.032 | **-0.819\*\*\*** | 0.037 | **1.356\*\*\*** | 0.834 | **-0.829\*\*\*** | 0.562 | **1.647\*\*\*** | -0.58 | 0.004 |
| BMI | 0.244 | 0.06 | **-0.537\*\*** | -0.066 | **1.053\*\*\*** | 0.515 | **-0.519\*\*** | -1.003 | 0.534 | 1.034 | 0.013 |
| Quetelet Index | -0.058 | 0.003 | -0.264 | 0.088 | 0.285 | 0.042 | -0.264 | 0.047 | 0.263 | 0.045 | 0 |
| Leg Length | 0.06 | 0.004 | -0.258 | -0.059 | 0.426 | 0.088 | -0.257 | -0.093 | 0.408 | 0.037 | 0 |
| CMJ | -0.138 | 0.019 | **-0.524\*** | 0.134 | **0.531\*** | 0.14 | **-0.497\*** | -1.333 | -0.28 | 1.618 | 0.031 |
| SJL | 0.055 | 0.003 | **-0.336\*** | 0.003 | **0.530\*** | 0.14 | -0.332 | -0.215 | 0.409 | 0.24 | 0.031 |
| SJR | 0.124 | 0.015 | **-0.341\*\*** | 0.018 | **0.631\*\*** | 0.181 | -0.354 | 0.737 | 1.029 | -0.793 | 0.007 |
| STJ | 0.028 | 0.001 | 0.049 | -0.017 | -0.105 | 0.005 | 0.039 | 0.557 | 0.212 | -0.633 | 0.005 |
| IAT | 0.077 | 0.006 | -0.062 | 0.151 | -0.01 | 0.023 | -0.091 | 1.734 | 0.866 | -1.746 | 0.036 |
| 505 Agility Test | 0.195 | 0.038 | 0.067 | 0.048 | -0.351 | 0.06 | 0.032 | 1.958 | 0.706 | -2.107 | 0.052 |
| 15-m Sprint | 0.162 | 0.026 | 0.288 | 0.02 | **-0.608\*** | 0.17 | 0.261 | 1.493 | 0.207 | -1.625 | 0.031 |
| 30-m Sprint | 0.042 | 0.002 | 0.125 | -0.062 | -0.117 | 0.009 | 0.094 | 1.639 | 0.825 | -1.877 | 0.042 |
| 5×25 Sprints | 0.291 | 0.085 | -0.012 | -0.054 | 0.407 | 0.080 | 0.006 | -1.025 | -0.130 | 1.070 | 0.014 |
| Spot Passing | -0.203 | 0.041 | -0.161 | 0.040 | -0.054 | 0.003 | -0.157 | -0.150 | -0.159 | 0.209 | 0.001 |
| Shooting | -0.079 | 0.006 | -0.053 | 0.039 | -0.033 | 0.002 | -0.045 | -0.391 | -0.271 | 0.474 | 0.003 |
| Dribbling | -0.245 | 0.060 | -0.112 | -0.091 | -0.186 | 0.022 | -0.160 | **2.524\*** | 1.260 | **-2.883\*** | 0.098 |
| AM-S | 0.161 | 0.026 | 0.242 | -0.108 | -0.118 | 0.017 | 0.236 | 0.222 | 0.064 | -0.364 | 0.002 |
| AM-F | -0.240 | 0.058 | -0.230 | 0.030 | -0.012 | 0.001 | -0.250 | 1.139 | 0.602 | -1.223 | 0.018 |
| AM | -0.252 | 0.063 | -0.100 | 0.175 | -0.194 | 0.051 | -0.122 | 1.382 | 0.474 | -1.331 | 0.021 |
| T | -0.096 | 0.009 | -0.166 | 0.102 | 0.101 | 0.014 | -0.215 | **2.781\*** | 1.583* | **-2.955\*** | 0.103 |
| E | -0.190 | 0.036 | -0.145 | 0.151 | -0.051 | 0.025 | -0.183 | 2.266 | 1.119 | -2.333 | 0.064 |
| TEOSQ | -0.180 | 0.032 | 0.112 | 0.115 | -0.388 | 0.086 | 0.095 | 1.049 | 0.129 | -1.030 | 0.013 |
| F1 | 0.038 | 0.001 | 0.022 | -0.158 | 0.011 | 0.025 | 0.028 | -0.502 | -0.179 | 0.379 | 0.002 |
| F2 | -0.050 | 0.003 | 0.040 | 0.172 | -0.111 | 0.037 | 0.049 | -0.313 | -0.379 | 0.535 | 0.003 |
| F3 | -0.206 | 0.043 | -0.063 | 0.056 | -0.190 | 0.020 | -0.061 | -0.089 | -0.270 | 0.160 | 0.000 |
| F4 | -0.120 | 0.014 | -0.216 | 0.209 | 0.145 | 0.050 | -0.219 | 0.368 | 0.233 | -0.175 | 0.000 |
| F5 | -0.153 | 0.024 | -0.061 | 0.119 | -0.117 | 0.022 | -0.062 | 0.159 | -0.094 | -0.044 | 0.000 |

\*, $p < 0.05$ ; \*\*, $p < 0.01$ ; \*\*\*, $p < 0.001$ .
$\beta 1$ =Standardized Coefficients of Age, $\beta 2$ =Standardized Coefficients of Relative Age, $\beta 3$ =Standardized Coefficients of Biological Maturation, $\beta 4$ =Standardized Coefficients of Relative Age × Biological Maturation, CMJ=Countermovement Jump, SJL=Single-leg Jump (Left), SJR=Single-leg Jump (Right), STJ=Standing Triple Jump, IAT=Illinois Agility Test, AM-S=Achievement Motivation (Approach Success), AM-F=Achievement Motivation (Avoidance Failure), AM=Achievement Motivation, T=Task Orientation, E=Ego Orientation, TEOSQ=Task and Ego Orientation in Sports Questionnaire, F1=Goal-Oriented, F2=Emotional Control, F3=Positive Cognition, F4=Family Support, F5=Interpersonal Assistants, F=Youth Psychological Resilience Scale.

### 3.3 Hierarchical Regression Analysis

A hierarchical regression analysis was conducted to examine the main effects and interaction effects of biological maturation and RA. The models were structured as follows: Model 1. chronological age; Model 2. main effects of biological maturation and RA; and Model 3. interaction effects of biological maturation and RA.

Biological maturation played a dominant role in the physical development of U16 male elite soccer players in Shanghai. Among elite soccer players of the same chronological age, those with advanced biological maturation exhibited better physical performance. Compared to Model 1, Model 2 includes RA and biological maturation. Model 2 showed a significant increase in explanatory power for most anthropometr ic indicators, except for leg length. Both maturity and chronological age had a significant effect on most anthropometric indicators (*p* < 0.05), whereas RA did not show a significant effect. Model 3, which examined the interaction effect between biological maturation and RA, revealed that biological maturation still had a significant effect on height, body weight, and sitting height, but the increase in model explanatory power was not significant. The interaction effect between biological maturation and RA was not statistically significant (*p* > 0.05). The explanatory power of the physiological index was much lower than that of anthropometric index, which may be due to the long-term systematic soccer training leading to a certain homogenization in physical performance among this group. Biological maturation significantly influenced SJ (β = 0.531, 0.530, *p* < 0.05), STJ (β = 0.631, *p* < 0.05), and 30-m sprint (β = -0.608, *p* < 0.05). For psychology, most were not significantly influenced by biological maturation or RA, except for the achievement motivation (β = 2.524, *p* < 0.05) and ego orientation (β = 2.781, *p* < 0.05).

## 4 Discussion

### 4.1 The Main Effects of Biological Maturation in Anthropometry and Physiology

Early maturing players gain anthropometric advantages during early adolescence through earlier skeletal and muscle growth. Biological maturity significantly influenced performance in SJ (*p* < 0.05), STJ (*p* < 0.05), and 30-meter sprint (*p* < 0.05), consistent with findings from Baxter-Jones et al. ^[27]^. In contrast, RA showed no statistical significance in either anthropometric or physiological indicators. Johnson et al. ^[5]^ argued that from adolescence onward, early maturing players exhibit significant advantages in body composition and motor capabilities, likely due to increased secretion of testosterone and growth hormone during their earlier onset of puberty ^[24]^. Early maturing players are typically taller, heavier, faster, stronger, and more powerful than their later maturing peers of the same chronological age ^[4],[25]^. Given the significant impact of biological maturation on anthropometric or physiological performance, early maturing players may demonstrate superior results in competitive matches and physical tests. However, late maturing players with soccer talent may struggle to compete against physically superior peers, leading to their marginalization within soccer development systems. From a long-term perspective, an overemphasis on early maturing players may yield negative consequences, as these players may prioritize physical advantages over the cultivation of technical, tactical, and psychological ^[26]^. While short-term success may be achieved, this approach hinders long-term development, as maturity-related anthropometric or physiological differences typically diminish or reverse after adulthood. Late maturing players, facing limited access to high-quality training resources during their developmental phase. Therefore, it will be detrimental to the construction of the soccer talent system.

### 4.2 Relative Age Effect is the accumulated training load effect

This study found that biological maturation and RA had no significant impact on soccer-specific skills and psychology, which is consistent with the findings of Malina et al. ^[1]^. Biological maturity showed a significant positive correlation with passing (r = 0.403), likely due to the enhanced lower limb explosive power in early maturing players, which may influence long-distance passing. The absence of significant effects of RA on anthropometric and physiological indicators in this study can be explained by the nature of RAE, which primarily reflects systemic biases in talent identification based on accumulated training rather than actual physical development. Unlike biological maturation, which is driven by internal physiological mechanisms and directly influences performance test outcomes. The RAE did not alter a player’s inherent physical capabilities. Instead, it stemmed from the temporal differences introduced by age grouping (maximum 12 months), which may primarily influence players’ experience and subsequent development of technical and psychological skills. Previous studies indicated that the RAE mainly affected access to training opportunities, competitive experience, and coaches’ subjective judgments, rather than test results ^[2],[6],[7]^. Therefore, the RAE primarily influenced early-stage talent identification and selection processes, rather than directly impacting test performance.

### 4.3 Independent Mechanisms of Biological Maturation and RA in Soccer Talent Selection

This study found that RA and its interaction with biological maturation did not show significant effects on anthropometric or physiological indicators (*p* > 0.05). Players born early accumulated more experience and demonstrated higher cognitive levels compared to players born late. Biological maturation and RA exert independent effects on the talent selection criteria of U16 elite soccer players, rather than biological causing the RAE. The differences induced by biological maturation emerged as puberty ^[1], [4], [5]^. RAE persists from late childhood to adolescence, while biological maturation significantly strengthens during adolescence and then diminishes after the end of adolescence. RA is the strongest influencing factor in soccer talent selection during childhood, while biological maturation becomes more critical during adolescence ^[10]^.

Talent identification must clearly distinguish between the selection biases caused by biological maturation and RA, and efforts to reduce the RAE should focus on dimensions more closely related to age, such as psychology, soccer-specific skills, and experience, and should be implemented from grassroots levels. Strategies for addressing selection biases related to biological maturation are more appropriate during adolescence, as differences in this period are more pronounced. It is important to recognize the independence of biological maturation and RA, as it is possible within the same age group to have players who are the oldest in chronological age but the least mature, and vice versa.

## 5 Limitations and Future Directions

This study is a cross-sectional study which limits its ability to track the dynamic relationship between biological maturity and performance indicators and thus cannot analyze the changes in these indicators during different stages of adolescence (before/after APHV). By increasing the diversity of the sample, a more comprehensive understanding of the influence of biological maturity on player selection in various contexts can be achieved, thereby enhancing the reliability and generalizability of the findings. In terms of biological maturity assessment, this study primarily relied on prediction equations based on anthropometric indicators. However, existing formulas are associated with certain inaccuracies. Future study could explore more precise methods for evaluating biological maturity or adopt costlier radiographic skeletal age assessment to improve the accuracy of maturity evaluation, thus providing a more reliable basis for talent selection.

## 6 Conclusion

Biological maturation significantly influences anthropometry (height, weight and Quetelet index), lower limb explosive, and speed (single-leg jump, standing triple jump, and 30-m sprint) in U16 male elite soccer players in Shanghai. The REA shows no significant impact on talent selection indicators, which is attributed to the accumulated training load effect. The mechanisms through which biological maturation and RA influence the selection of young soccer players are distinct, exhibiting relatively independent roles.

## Data available

All the data generated or analyses during this study are included in this article. This article received no funding.

